# Dynamic expression of smRNA from fecal exosome in disease progression of an inflammatory bowel disorder mouse model

**DOI:** 10.1101/2021.02.24.432648

**Authors:** Sean Manning, Shisui Torii, Hannah M Atkins, Yuka Imamura Kawasawa

## Abstract

Inflammatory bowel disease (IBD) is a chronic inflammatory disorder of the gastrointestinal tract affecting over 3 million adults in the United States. Despite being widespread, reliable early diagnostic tests are not available. We examined exosomal small RNA (smRNA), specifically targeting microRNA (miRNA) and piRNA from the stool samples of IBD model mice, interleukin 10 knockout mice (IL-10 KO), as a potential diagnostic marker. Stool samples were specifically chosen because they are readily available, and collection is noninvasive. At the end of the experimental period, the gastrointestinal (GI) tract was collected, and disease severity was scored. Histopathology showed a significant increase in inflammation and proliferation within the proximal and distal large intestines. smRNA profiles were examined upon conventional housing (start-point) which is a determinant factor of spontaneous IBD progression in the IL-10 KO mice, terminal illness (end-point), and 6 weeks before the end-point (mid-point), when the mice were still phenotypically healthy. We found 504 smRNA that were significantly differentially expressed between before symptom onset and terminal sedation. These changes were not detected in wild-type samples. Moreover, clustering analysis of expression changes over the disease progression identified a unique set of smRNAs that primarily target pro-inflammatory or anti-inflammatory genes. The expression of smRNAs that suppresses pro-inflammatory genes was higher at 6 weeks before terminal sedation, suggesting the downregulation of the pro-inflammatory genes advances the terminal illness of the IBD. In summary, our study proposes that fecal exosomal smRNA profiling offers a new opportunity to monitor the inflammatory status of the gut with a capability of detecting its pro-inflammatory (asymptomatic) status. Our next step is to understand the spatiotemporal interplay of these exosomes and the host cells in the gut as well as the complete biochemical makeup of fecal exosomes, such as mRNA, DNA, protein, and lipids. This will lead to an exciting development of reengineered exosomes that can be utilized to treat or even prevent the pro-inflammatory colonic lesion while the host is still clinically asymptomatic.

## INTRODUCTION

Inflammatory bowel disease (IBD), commonly categorized into either Crohn’ s disease (CD) or ulcerative colitis (UC), is a common cause of chronic inflammation in the gastrointestinal tract (1). While often referred to under the same umbrella term IBD, CD and UC have several differences. Patients with CD often have patches of disease through the whole gastrointestinal (GI) tract, with inflammation affecting the full thickness of the intestine. In contrast, UC manifests mostly in the colon and rectum with inflammation often limited to the mucosa (2). Clinical signs usually consisting of diarrhea, fever, and abdominal pain are often cyclic, with flare-ups followed by remission periods (3). In all cases, IBD leads to a low quality of life, often with multiple hospitalizations and surgical procedures (4).

While over 3 million adults in the US are affected, diagnosis is often prolonged (5). According to the European Crohn’ s and Colitis Organization [ECCO], “a single reference standard for the diagnosis of Crohn’ s disease [CD] or ulcerative colitis [UC] does not exist” (6). Current methods for diagnosing IBD include a combination of physical exams, blood tests, stool examinations, endoscopy, biopsies, and imaging (7). This battery of tests results in an average 2-3-month delay in diagnosis for UC and 6-7-month delay for CD (8, 9). More recently, the study of exosomes and the exosomal small RNA (smRNA) they contain have shown to be a valuable diagnostic marker in IBD and many other diseases.

Exosomes are a sub-type of extracellular nano-sized vesicle that have been shown to mediate extracellular communication via the transport of various biomolecules. Exosomes transport coding and non-coding RNA, including smRNA. This signaling process impacts several physiological and pathological pathways by modulation of protein expression. Due to their involvement in critical biological functions, studies into the use of exosomal smRNA as biomarkers in various diseases and tissues have grown significantly (10–12). For example, serum miRNA show promise as a diagnostic test, differentiating colitis subtypes, and evaluating therapy response (13). While studied extensively in serum, intestinal biopsies, and even saliva (2, 13), little is known about fecal exosomes’ role in disease development. In contrast to serum, feces are abundant, GI tract-specific, and noninvasive to collect. Studies such as Wall et al. (14) investigating fecal exosomes in colorectal cancer screening, and Liu et al. (15) investigating fecal miRNA and the microbiome, show value in the use of feces to evaluate miRNA. However, fecal exosomes in IBD have not been studied.

Our study evaluated fecal smRNA profiles and tracked changes before and after IBD development in IL-10 KO mice and compared them to wild-type mice. IL-10 KO mice have been extensively studied as an IBD model (16–20). Although IL-10 KO mice are viable and fertile when housed under specific pathogen free (SPF) conditions, once they are conventionally housed, they develop altered lymphocyte and myeloid profiles, elevated serum amyloid A levels, altered responses to inflammatory or autoimmune stimuli, increased prevalence of colorectal adenocarcinoma, and spontaneous development of chronic enterocolitis (The Jackson Laboratory Cited 25 March 2018). While spontaneous enterocolitis is the target of this study, human IBD patients can also similarly develop secondary systemic amyloidosis (SSA) (21) are prone to the development of colorectal adenocarcinoma (22), and have dysfunctional immune responses (1). We hypothesize that fecal exosomal smRNA shows a unique profile in disease progression profile in the IL-10 KO mice. This unique fingerprint is a promising, noninvasive diagnostic tool for IBD.

## METHODS

### Animals

Four-week-old IL-10 KO mice (Stock No. 002251 with C57BL/6J background, IMSR Cat# JAX:002251, RRID:IMSR_JAX:002251) and control wild-type (WT) mice (C57BL/6J, MGI Cat# 5657312, RRID:MGI:5657312, equal number of females and males was used) were obtained from The Jackson Laboratories (Bar Harbor, Maine). Each mouse was then housed conventionally and individually in polycarbonate cages, static, wire-top cages on corncob bedding (7092 Harlan Teklad, Madison, WI), and had *ad lib* access to irradiated rodent chow (2918, Harlan Teklad, Madison, WI) and tap water. The animal facility was programmed with a 12:12-h light: dark cycle. Room temperature was maintained at 20 ± 2°C, and air humidity ranged between 30 to 60%. Environmental enrichment was provided in the form of an Enviropak (Lab Supply, Fort Worth, TX). Mice were free of ectromelia virus, Theiler’ s encephalomyelitis virus, Hantaan virus, K virus, lactate dehydrogenase elevating virus, lymphocytic choriomeningitis virus, mouse adenovirus types 1 and 2, mouse cytomegalovirus, mouse hepatitis virus, mouse minute virus, murine norovirus, mouse parvovirus, mouse thymic virus, pneumonia virus of mice, polyoma virus, Reovirus 3, Rotavirus, encephalomyelitis virus, Sendai virus, *Bordetella bronchiseptica, Flavum rodentium, Citrobacter rodentium, Clostridium piliforme, Corynebacterium kutscheri Corynebacterium bovis, Helicobacter spp*., *Mycoplasma pulmonis, Pasteurella spp*., *Salmonella spp*., *Streptobacillus moniliformis, Klebsiella pneumonia, Pneumocystis murina, Pseudomonas spp*., *Staphylococcus aureus, Streptococcus pneumonia*, β-hemolytic *Streptococcus spp*., endo-and ectoparasites, enteric protozoa, and *Encephalitozoon cuniculi*. All experiments were conducted in accordance with institutional guidelines, the Guide for the Care and Use of Laboratory Animals (Institute for Laboratory Animal Research, 2011), and approved by the Penn State College of Medicine Institutional Animal Care and Use Committee.

### Fecal Collection

The feces were collected once a week and stored at -80°C until processing. Mice were scuffed, and a 1.5 mL Eppendorf tube (Fisher Scientific, Waltham, MA) was placed under the anus to collect 3-5 fecal pellets. If sufficient feces were not collected after 30 seconds, the mouse was placed in a clean cage without beading until at least 3 pellets were obtained. During this time, the mice were also scored for disease severity using a modified Crohn’ s Disease Activity Index (CDAI) based on Scheinin et al. (20) (Table 1). Collection continued until mice either reached a score of 3 for three consecutive times or a score of 4, which were established as experiment end-points. The fecal samples collected at the first week of conventional housing and 6 weeks before the experimental end-point were designated as start-point samples and mid-point samples, respectively (Figure 1). Some KO mice never developed IBD (scored 0 (no signs of disease) for the entire experimental period) and were euthanized after 30 weeks of conventional housing (Figure 1). For WT mice (no signs of disease throughout the 30 weeks of conventional housing), we collected the feces samples starting at the average age of disease onset in KO mice (5 months) for a duration of 6 weeks. At the time of euthanasia, the gastrointestinal tract was collected. The small and large intestines were separated just proximal to the cecum and fixed in 10% neutral buffered formalin for 24 hours before being transferred to 70% ethanol. The tissues were paraffin-embedded using an automated embedding machine and sectioned 5-um thick for routine Harris hematoxylin and eosin (HE) staining. The intestines were microscopically evaluated and scored by a board-certified veterinary pathologist blinded to the groups. Five randomly-selected, 20X fields of small or large intestine were scored for degree of hyperplasia, inflammation severity, goblet cell loss, and epithelial cell erosion or ulceration. Each microscopic characteristic was assigned a severity score from 0-4, based on Erben et al. (23) (Tables 2 and 3). The 20X fields were averaged and a total severity score was calculated for each intestine region (proximal and distal small intestine, cecum, and proximal and distal large intestine).

**Fig 1.**
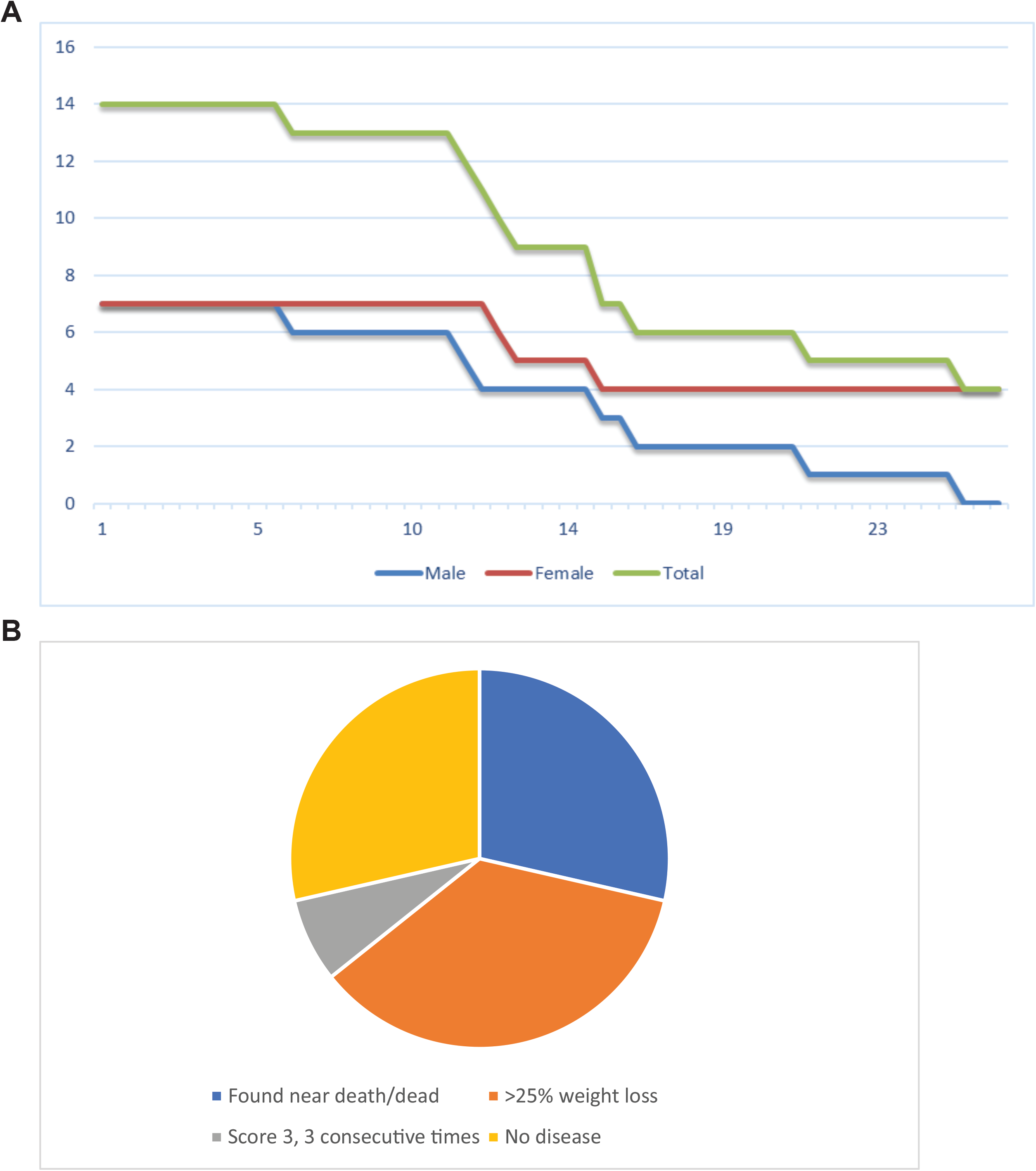
Survival curves for IL-10 KO mice. (A) The number of survivors at the indicated weeks is shown for IL-10 KO. The number of mice euthanized before achieving humane endpoints was counted. (B) The pie chart shows the proportion of the cause of death in IL-10 KO mice.

### Exosomal RNA Extraction

Exosomal RNA was extracted and sequenced as follows from start-point (S), mid-point (M; 6 weeks prior to the experimental end-point), and experimental end-point (E) that is defined as either when the animals reached a score of 3 for three consecutive times or a score of 4, or after 30 weeks if the animals did not develop clinical disease (Figure 1). We developed a novel method to reliably extract exosomal RNA from mouse feces. Briefly, 2 fecal pellets (∼60mg) were vortexed with 0.9-2.0 mm stainless steel beads (Next Advance, Troy, NY) with 60 μl Tris-HCl (10 mM) in a 1.5 mL microcentrifuge tube. Increasing amounts of Tris-HCl until 1 mL total volume was reached and subsequently vortexed. The resulting mixture was then centrifuged at 10,000Xg for 5 minutes. The supernatant was then filtered through a 0.2 μm syringe filter (ThermoFisher Scientific, Waltham, MA). Exosomes were isolated via ExoQuick TC (System Biosciences, Palo Alto, CA) according to manufacturer instructions. For quality check of exosomes, a transmission electron microscope (TEM, JEOL-1400 transmission electron microscope (JEOL USA, Peabody, MA)) and a particle tracking system, NanoSight (NanoSight Ltd, Salisbury, UK) were used. The remainder of the purified exosomes were lysed and RNA was isolated using Quick-RNA Microprep Kit (Zymo Research, Irvine, CA). The quality and quantity of the extracted RNA were determined using TapeStation RNA ScreenTape (Agilent Technologies, Santa Clara, CA) and Qubit fluorometer (ThermoFisher Scientific).

### Small RNA sequencing and sample quality assessment

SmRNA-sequencing libraries were prepared from 50 to 150 ng of total RNA using the QIAseq miRNA Library Kit (Qiagen, Germantown, MD) as per the manufacturer’ s instructions. This system offers a built-in Unique Molecular Identifier (UMI) application, which is used to eliminate possible PCR duplicates in sequencing datasets and therefore facilitate unbiased gene expression profiling. The unique barcode sequences were incorporated in the adaptors for multiplexed high-throughput sequencing. The final product was assessed for its size distribution and concentration using BioAnalyzer High Sensitivity DNA Kit (Agilent Technologies). Pooled libraries were diluted to 2 nM in EB buffer (Qiagen) and then denatured using the Illumina protocol. The denatured libraries were loaded onto an SP flow cell on an Illumina NovaSeq 6000 and run for 53-71 cycles using a single-read recipe according to the manufacturer’ s instructions. De-multiplexed sequencing reads were generated using Illumina bcl2fastq (released version 2.20.0.422, Illumina), allowing no mismatches in the index read. Primary read mapping and UMI analysis were conducted via the GeneGlobe Data Analysis Center (Qiagen).

### Differential expression analysis of smRNA

Input raw read count data was used after eliminating the genes for which counts per million (CPM) of reads were lower than 1 on average. edgeR R package (RRID:SCR_012802) was used to normalize the raw read counts to obtain Trimmed Means of M (TMM) values. Integrative Differential Expression Analysis for Multiple EXperiments (IDEAMEX) (24) was used for four alternative differential expression analyses, namely, edgeR, DESeq2 (RRID:SCR_015687), NOI-seq (RRID:SCR_003002), and limma-voom (RRID:SCR_010943). A multidimensional scaling (MDS) plot, a Venn diagram, and a volcano plot were also generated in IDEAMEX. An adjusted *p*-value of less than 0.05 and fold change cut off at 2 were used as a significance cutoff. ComplexHeatmap R package (RRID:SCR_017270) was used to create a heatmap. Short time-series expression miner (STEM, RRID:SCR_005016) was used for time-course analysis. Ingenuity Pathway Analysis (IPA, Qiagen, RRID:SCR_008653) was used to analyze functional enrichment in the differentially expressed smRNA. MicroRNA Target Filter function in IPA was also used to obtain the list of predicted target genes of these smRNAs. Enrichr (25) (RRID:SCR_001575) was used to examine functional enrichment of the predicted target genes.

### qRT-PCR

miRNA All-In-One cDNA Synthesis Kit (Applied Biological Materials, Richmond, BC, Canada) was used to convert RNA to generate. Real-time PCR amplification was performed using the PowerTrack SYBR Green Master Mix (ThermoFisher Scientific). 3 ng of input RNA was polyA-tailed, reverse transcribed, and PCR amplified using with a 5′-primer for mmu-miR-682 (CTGCAGTCACAGTGAAGTCTG) and the universal 3′ miRNA primer (Applied Biological Materials). Relative Ct values were calculated based on start-point samples and compared to the mid-(only for IL-KO) and end-point samples of the same mouse individuals. 6 animals were examined for each group (IL-10 KO and wild-type) and a one-tailed paired *t*-test was used to evaluate the statistical difference. A line plot was generated using gglot2 R package (RRID:SCR_014601).

### Statistical Analysis

Histopathology statistical analysis was completed using Tukey’ s multiple comparisons test and Sidak’ s multiple comparison test (GraphPad Prism8, San Diego, CA, RRID:SCR_002798) for both the small and large intestinal sections and comparisons between sexes. Statistical significance was defined as a *p-*value of less than or equal to 0.05.

## RESULTS

In IL-10 KO mice, all males (7/7, 100%) and only 3/7 (42.9%) of females were euthanized due to reaching experimental or humane endpoint (Figure 1A). No control mice were euthanized or perished by 30 weeks old (data not shown). IL-10 KO mice generally gained weight until disease onset, after which they declined. All control mice continually gained weight until the end of the experimental period. Cause of the death includes spontaneous death, significant loss of weight, or euthanization due to disease scoring more than 3 for three consecutive times. However, ∼30% of animals did not show any symptoms by 30 weeks old (Figure 1B).

During the necropsy of IL-10 KO mice, especially ones with high disease scores, had thickened colons with scant to no feces present compared to controls (Figure 2). The colons of IL-10 KO mice with no clinical signs appeared grossly normal for mice of their strain, sex, and age (Figure 2A). IL-10 KO mice showed a significantly higher score in the large intestine in both the proximal and distal sections (Figure 2B). There was no significant difference between disease severity in the small intestine or cecum sections (Figure 2B). For scores broken down by category in the distal large intestine, a significantly higher score was seen for IL-10 KO mice in inflammation location and proliferation (Figure 2B). For necrosis, severity trended towards significance; however, the sample size limited the analysis (Figure 2B). The same trend was also seen in the proximal large intestine (Figure 2B). No significant differences were seen between female and male scores for either the proximal or distal large intestine (Figure 2B). Representative images of the colonic histopathology are shown in Figure 2C.

**FIg 2.**
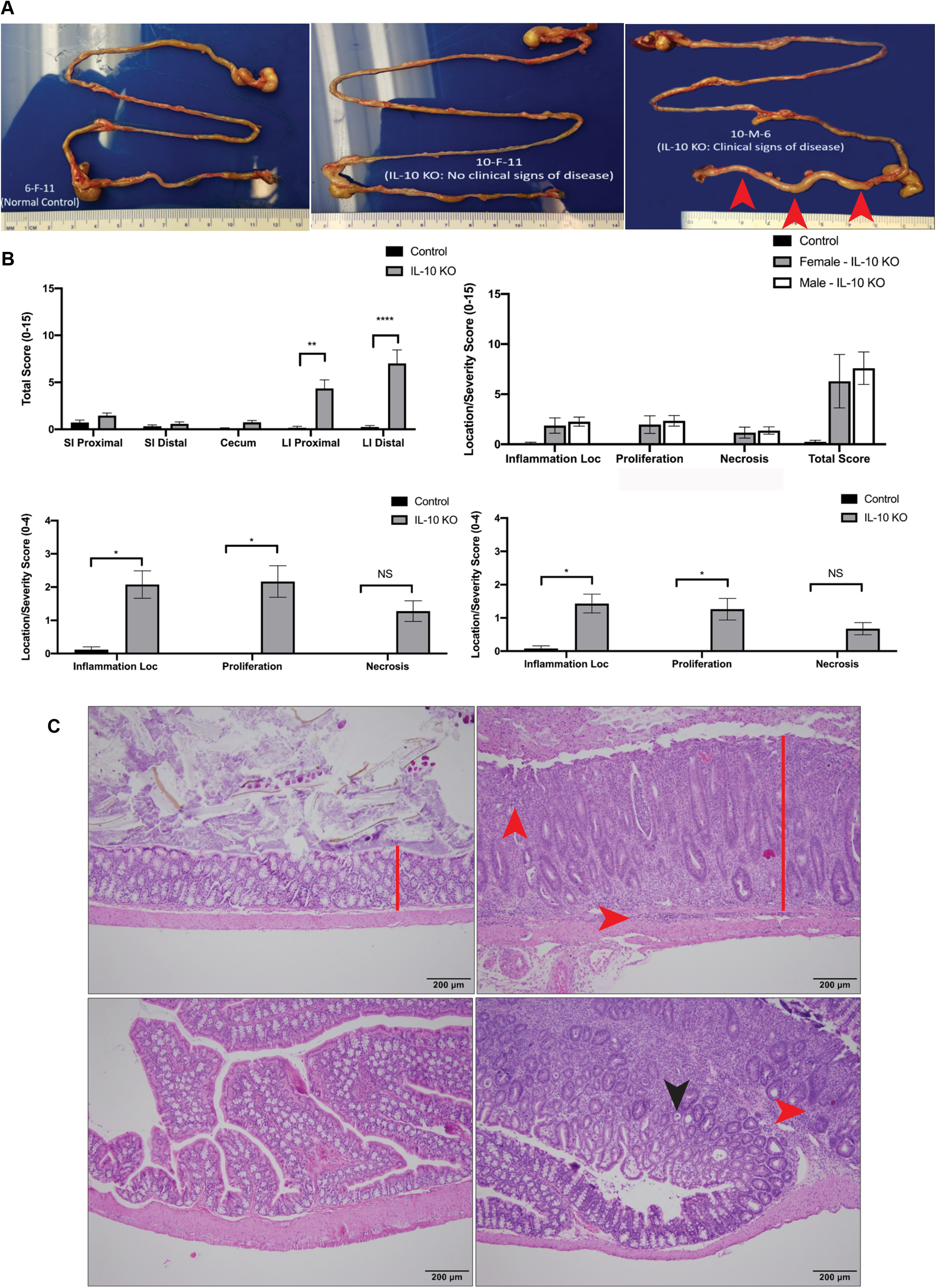
Pathology of the intestine in IL-10 KO mice. (**A**) Gastrointestinal tracts at necropsy from wild-type control (left), asymptomatic IL-10 KO (middle) and symptomatic (right) IL-10 KO mice. The large intestine from the symptomatic IL-10 KO mouse was thickened with scant to no fecal material present (red arrowhead). (B) (top left) Cumulative disease scores from four measures in indicated regions are compared between wild-type control and IL-10 KO mice. (top right) Cumulative disease scores and the scores of three measures are similar in female and male IL-10 KO that were sympomatic. Inflammation location is indicated as Inflammation Loc. (lower left and right) Comparison of the indicated measures in distal (left) and proximal (right) intestine between the indicated genotypes. *, **, and **** indicates *p*<0.05, <0.01, and <0.0001, respectively. NS indicates comparison which does not show significant difference. (C) Hematoxylin and eosin-stained wild-type control distal intestine sections (top left), clinically affected IL-10 KO (top right). Multifocal mucosal and submucosal infiltration of mixed inflammatory cells (red arrowheads) with epithelial hyperplasia (red line) and loss of goblet cells were found. Sections of the distal large intestine of wild-type control (lower left) and affected IL-10 KO (lower right). Multifocal infiltration of mixed inflammatory cells (red arrowhead) with crypt abscessation (black arrowhead). Scale bar = 200 μm.

Exosomes were verified using a JEOL-1400 transmission electron microscope (Figure 3A). Additionally, the sample was run through NanoSight analysis and determined 8.89 ×10^10^ +/-1.79 ×10^9^ particles/mL of exosomes in the sample ranging from 75-159 nm in size (Figure 3B). After RNA purification, TapeStation validated the presence of a large amount of smRNA (Figure 3C). Figure 4 also validated the quality of the smRNA-sequencing by confirming the similar levels of the gene detection between samples.

**Fig 3.**
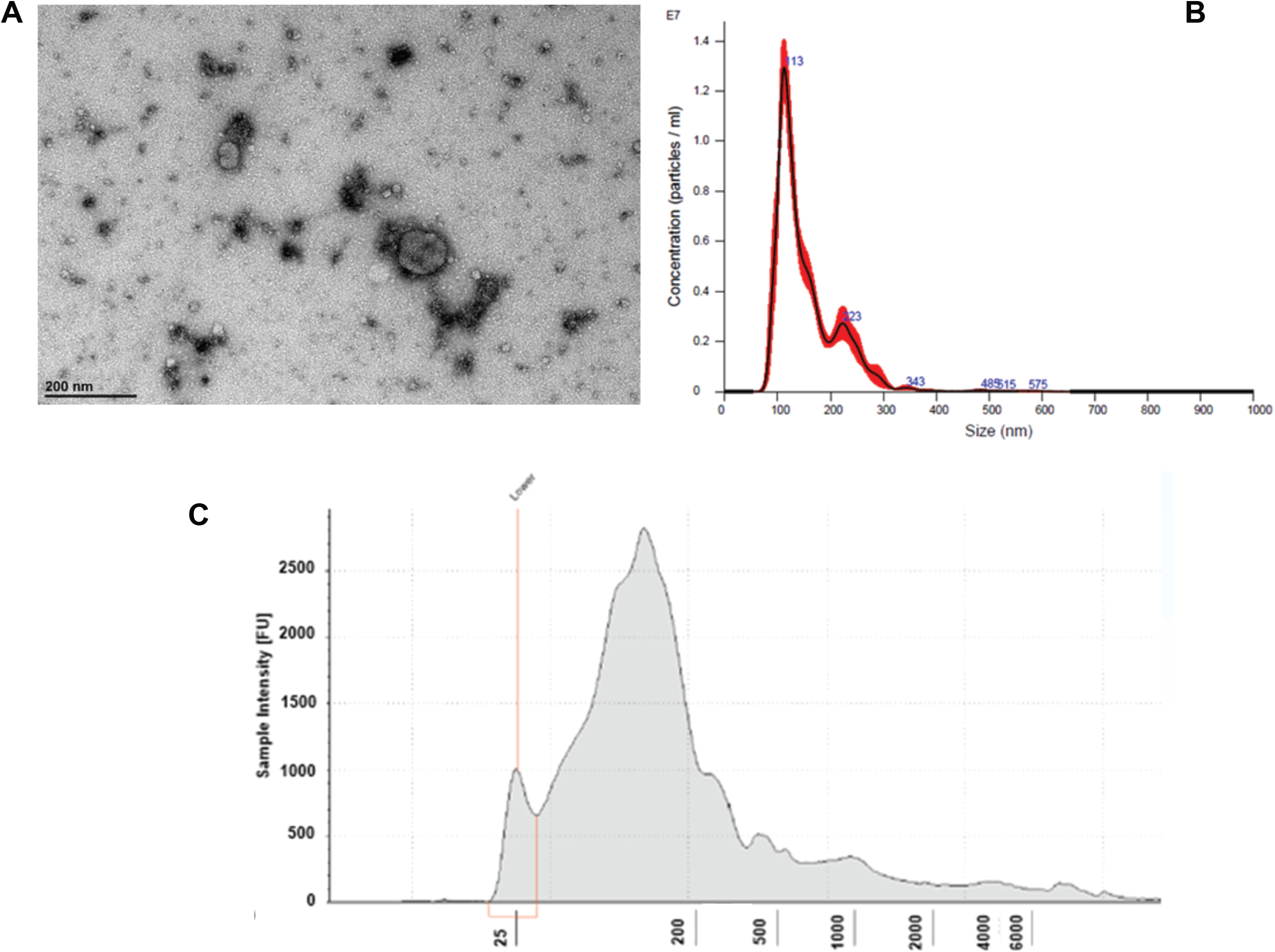
Extraction of exosomal smRNA from feces of mice. (A) transmission electron microscope image of exosomes taken from the feces of a wild-type mouse. (B) NanoSight analysis shows a large concentration of particles with ∼100 nm size, suggesting the presence of many exosomes. (C) TapeStation RNA ScreenTape shows the presence of a large amount of smRNAs.

**Fig 4.**
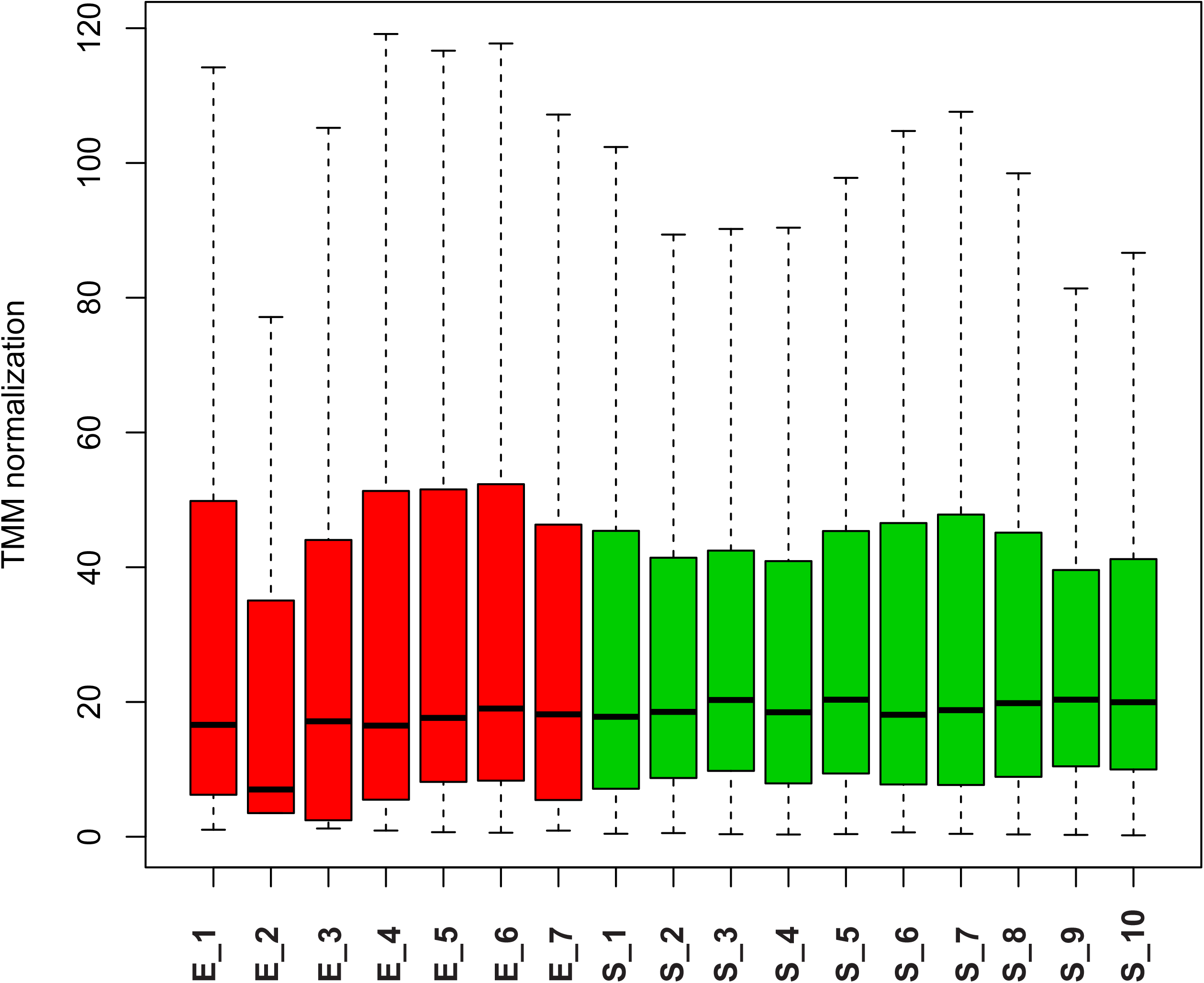
Quality control of smRNA sequencing. TMM normalization of RNA expression level for each sample.

The smRNA sequencing data of start-(S) and end-(E) points detected a total of 1,472 smRNAs that are expressed more than 1 CPM on average. Four female animals that didn’ t develop symptoms and samples that didn’ t pass our quality control were excluded from the analysis. The MDS plot shows that S and E samples manifest gross differences in smRNA profiles (Figure 5A). IDEAMEX offers a WEB-based platform to perform differential expression analyses of RNA-sequencing by four different R packages, edgeR, DESeq2, NOI-seq, and limma-voom, as well as integration of the information outcomes. The integrated Venn diagram of smRNAs that are defined as differentially expressed shows variable outcome, with only 40 smRNAs consistently called significantly different among 4 algorithms (Figure 5B and Supplementary Figure S1). We selected the outcome from edgeR analysis which has the largest number (504) of smRNA for subsequent analyses (Figure 5B, 5C and Supplementary Table S1). A heatmap is showing the top 50 differentially expressed smRNAs, with clear difference between S and E timepoints and no bias in the sex of the animals (Figure 5D).

**FIg 5.**
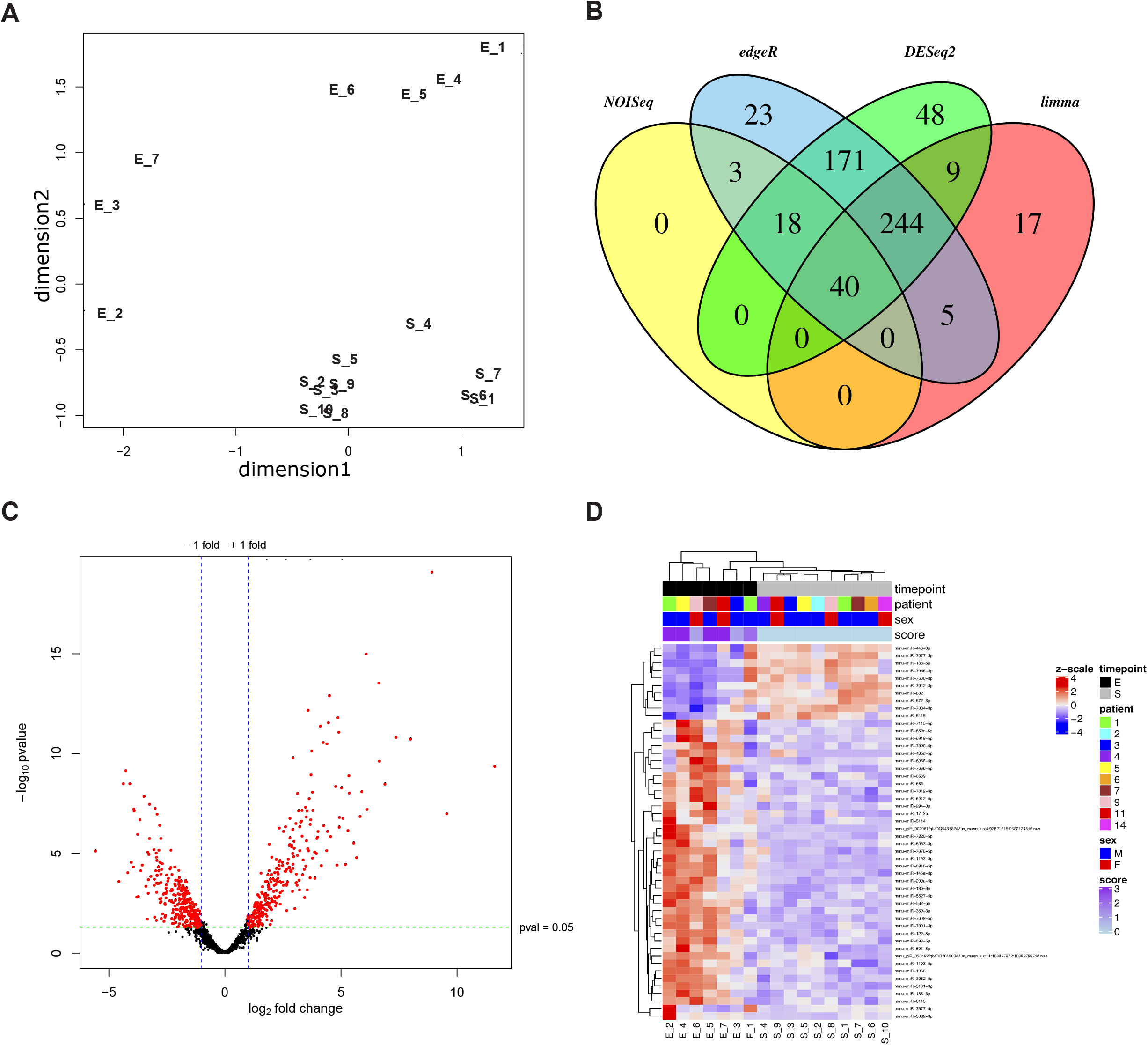
Differentially expressed genes between before symptom onset and terminal sedation. (A) MDS plot shows gross smRNA expression profiles are different between S (start-point) and E (end-point) samples. (B) Venn diagram of smRNAs that are defined as differentially expressed genes in edgeR, DESeq2, NOI-seq, and limma-voom. (C) The volcano plot shows the fold change and p-value of each smRNA. Red dots show genes which expressions are significantly different between S and E. (D) A clustered heatmap of the top 50 differentially expressed smRNAs is shown. The annotation bars depict sample information for its timepoint, mouse identity (patient), sex, and disease scores when feces were sampled, respectively.

Next, we performed the analysis to examine the change of expression throughout IBD development by using Short Time-Series Expression Miner (STEM) with the datasets at the start-, mid-, and end-point. Out of 16 clusters, 4 clusters include a significant number of the smRNAs than the predicted number based upon the entire number of smRNAs that were input into the STEM analysis (colored clusters in Figure 6A and Supplementary Table S1). Among the four significant clusters, the purple and pink clusters show sharp contrast. Expressions of smRNA in the purple cluster remain the same at the mid-point and decrease from mid-to end-point, while those in the pink cluster increase from start-to mid-point, and then decrease from mid-to end-point. We chose miR-682 from the pink cluster and examined its expression changes in wild-type and IL-10 KO mice by quantitative reverse transcription polymerase chain reaction (qRT-PCR). We were able to reproduce a significant reduction of miR-682 in the end-point samples while there was no change in wild-type. Mid-point samples in the KO mice also showed significant decrease (Figure 6B), which contradicts the upward trend in the STEM analysis (Figure 6A). Due to the limited amount of feces samples, we had to use RNA samples from different batches of preparations for the STEM analysis and qRT-PCR, which could be a reason for this variability.

**FIg 6.**
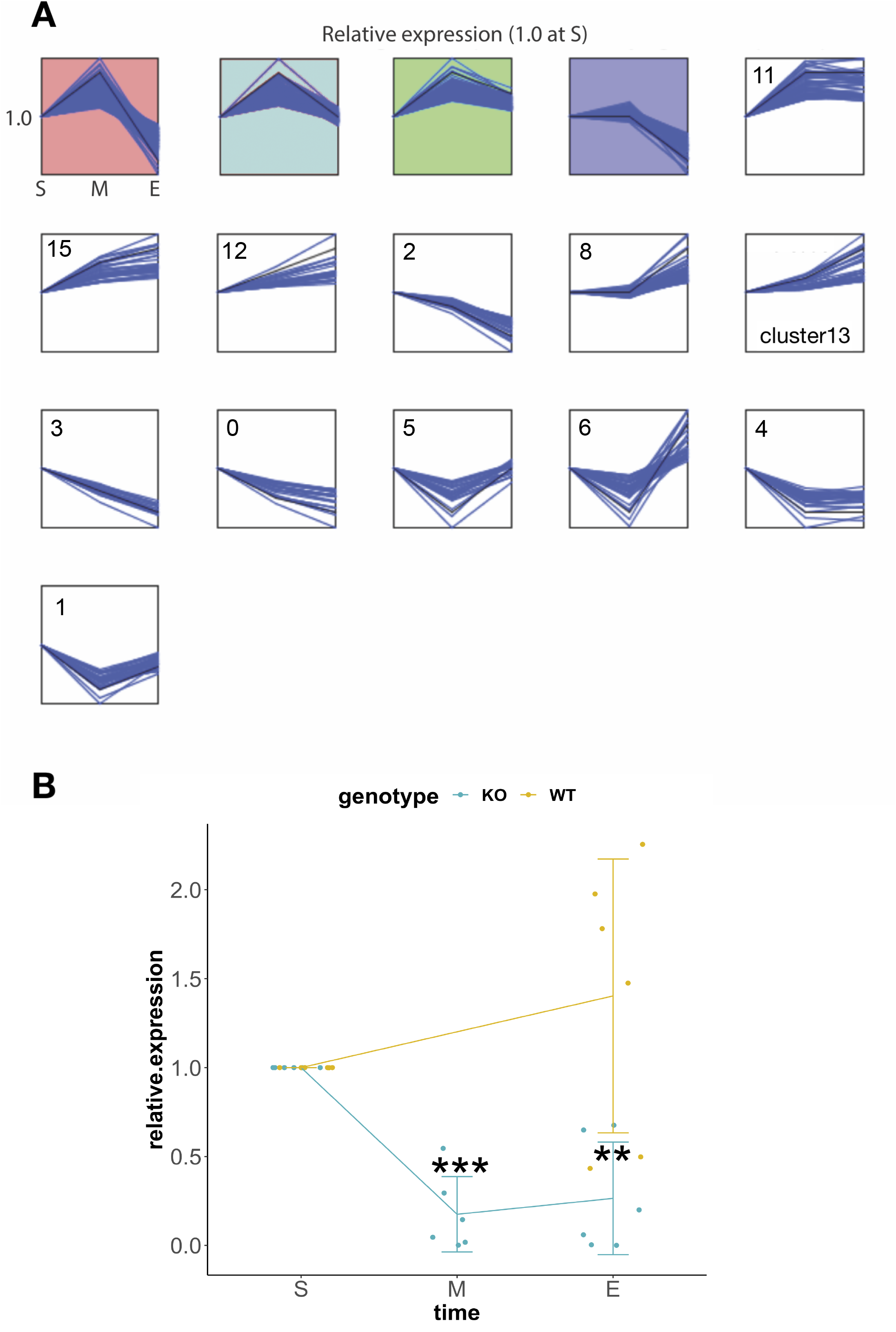
smRNA expression change pattern over the course of disease progression. (A) Colored clusters include a significant number of the smRNA molecules that show unique temporal changes. Blue and black lines indicate each smRNA and the average respectively. (B) Relative expression of mmu-miR-682 was validated by qRT-PCR and shown in the line plot. Error bars indicated the standard deviation of six animals chosen from both IL-10 KO (KO, light blue) and wild-type (WT). Fecal exosomal RNA was prepared from the same six mouse individuals at S (start-point), M (mid-point, KO animals only), and E (end-point). ** and *** indicate significant differences compared to S at *p*<<0.01 and <0.001, respectively.

To understand the functional relevance of these clusters, we performed functional annotation analysis of those two clusters. The purple cluster includes such as miR-208-5p, miR-367-5p, miR-342-3p, miR-4276, and miR-532-5p (Figure 7A). Target gene pathway analysis using the genes that are targeted by smRNA shows the enrichment of multiple pro-inflammatory pathways including the TNF-alpha, KRAS and IL-2/STA5 pathways (Figure 7B). The pink cluster includes three gene networks that are significantly regulated by the smRNA in this cluster, such as miR-19b-3p, miR-494-3p, miR-329-3p, miR-434-3p (Figure 8A). Both pro-inflammatory and anti-inflammatory pathways were found in the target gene pathway analysis (Figure 8B).

**Fig 7.**
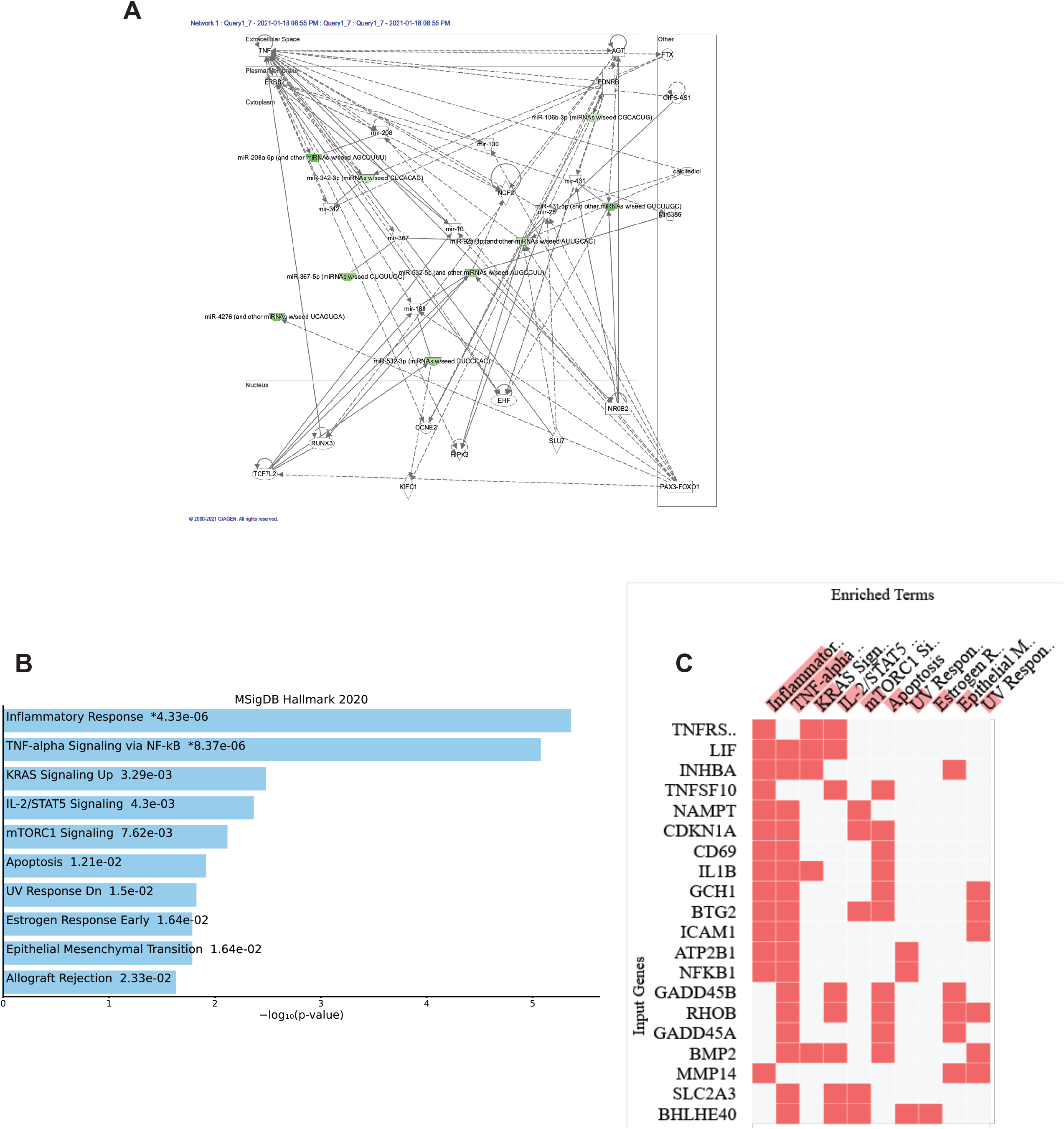
Major target genes of smRNA which expression is low at terminal sedation (purple cluster), including anti-inflammatory genes. (A) Gene network that is controlled by the down-regulated smRNAs (green) in the purple cluster. Cellular locations of the genes targeted by the smRNAs are also denoted. Dotted line and solid line indicate indirect and direct regulations, respectively. (B) Bar graph is showing significantly enriched functional pathway terms of the target genes of smRNAs in the purple cluster. Molecular Signatures Database (MSigDB) Hallmark version 2020 was used as a pathway dataset. (C) Clustergram of the genes that are included in the highly enriched functional pathways in (B). Anti-inflammatory genes are noticeably enriched.

**Fig 8.**
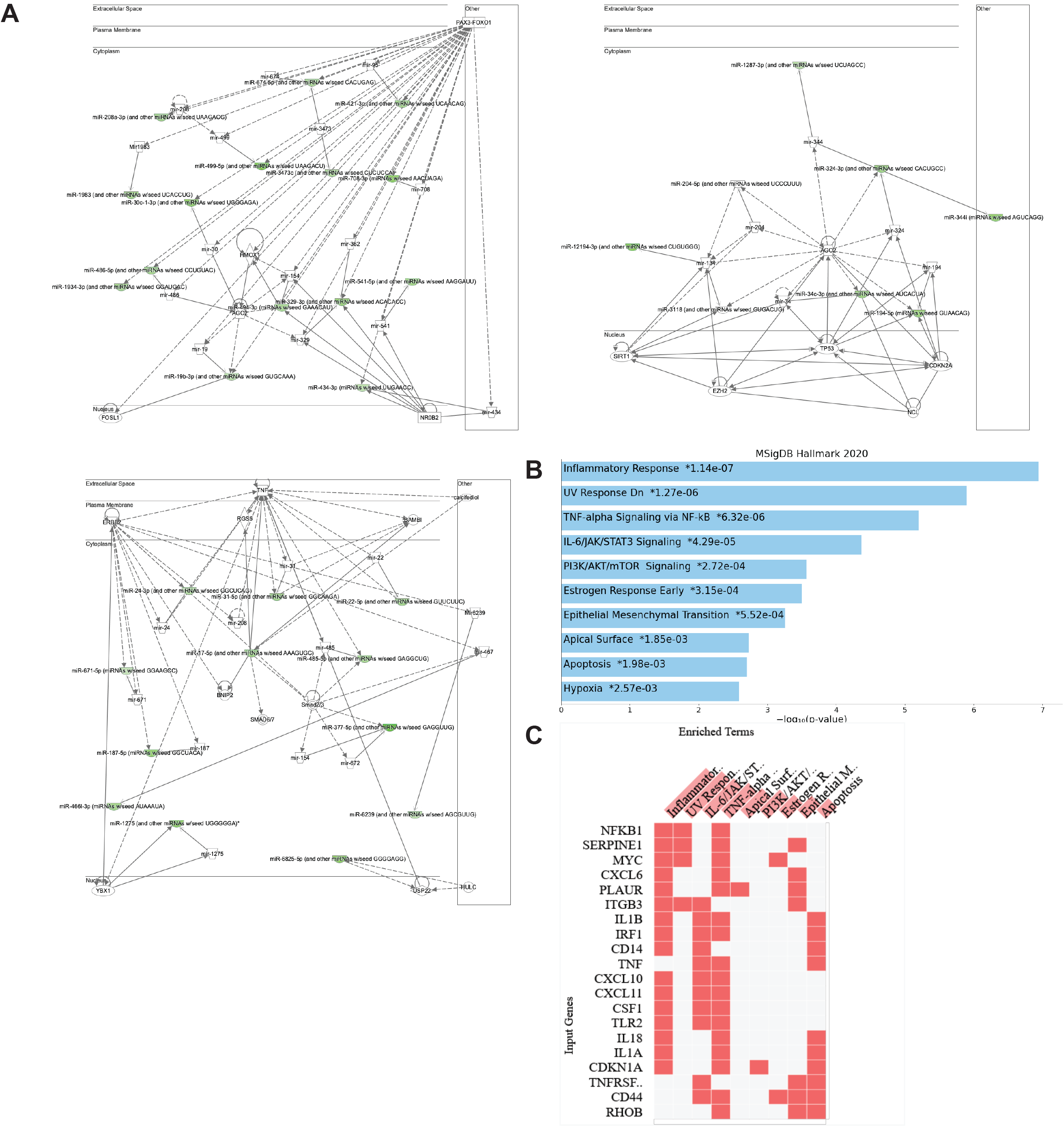
Major target genes of smRNA for which expression is high at 6 weeks before termination but low at terminal sedation (pink cluster), including pro-inflammatory genes. (A) Gene networks that are controlled by the down-regulated smRNAs (green) in the pink cluster. Three networks were predicted from the smRNAs in the pink cluster. (B) Bar graph is showing significantly enriched functional pathway terms of the target genes of smRNAs in the pink cluster. (C) Clustergram of the genes that are included in the highly enriched functional pathways in (B). Pro-inflammatory genes are enriched in this list.

Although the number of smRNA types is smaller, the cluster labelled as “cluster13” in Figure 6A and Supplementary Table S1 was further examined as this may work as the biomarkers of IBD progression (Figure 9). Although statistically non-significant, the smRNA in the “cluster13” shows the gradual increase of the expression compared with the other non-significant clusters that show a trend of the expression increase (clusters in the same row in Figure 6A). The IPA analysis did not yield a network that was controlled significantly by the smRNAs in this cluster, however, target gene pathway analysis revealed inclusion of unique pathways such as Notch signaling and the P53 pathway.

**Fig 9.**
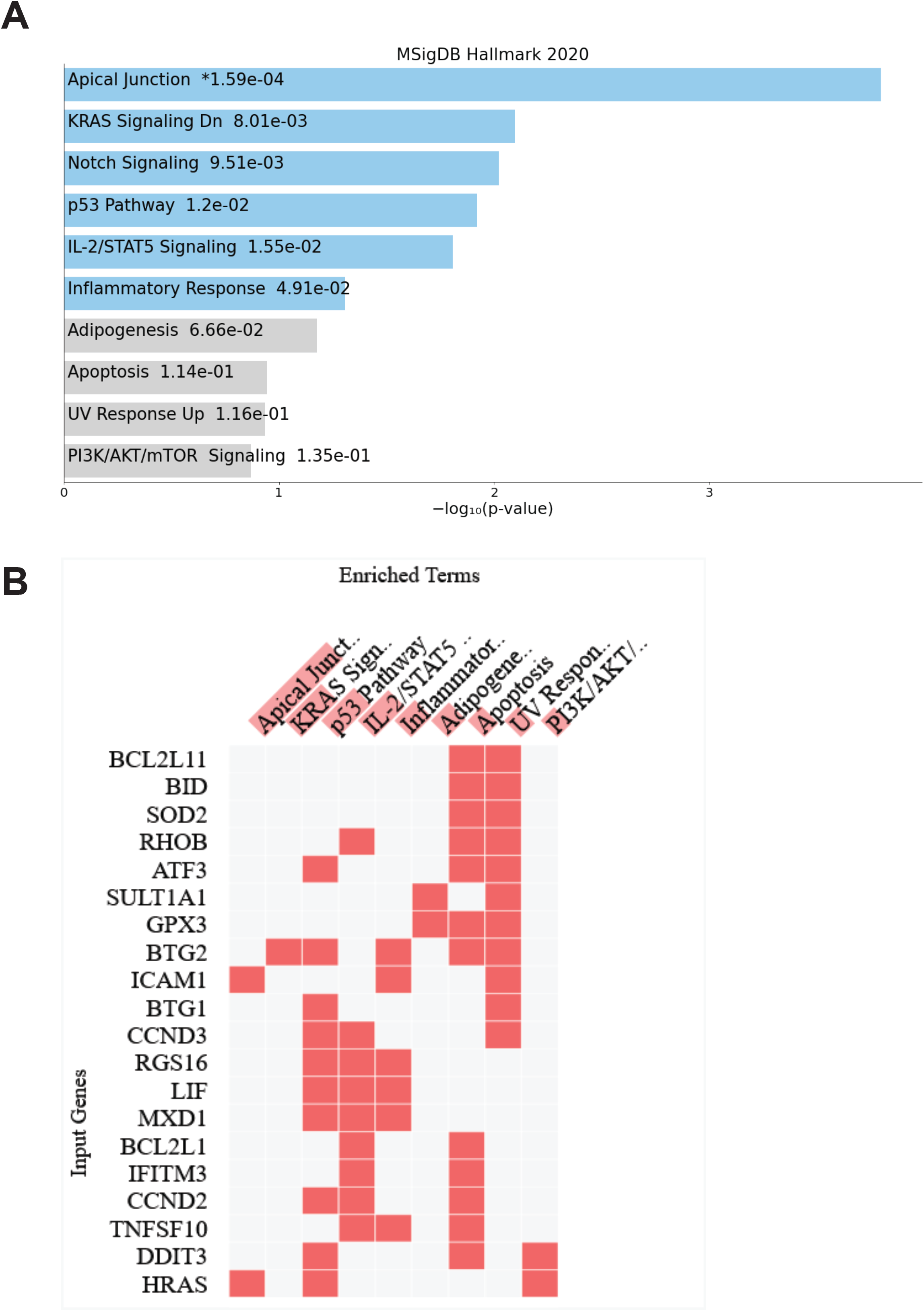
Major target genes of smRNA which expression increases over pathogenesis of IBD. (A) Bar graph is showing significantly enriched functional pathway terms of the target genes of smRNAs in the “cluster13” that has a consistent increase of the expression throughout pathogenesis. Statistically weak pathways are colored in gray. (B) Clustergram of the genes that are included in the highly enriched functional pathways in (A).

## DISCUSSION

Exosomes are a sub-type of extracellular nano-sized vesicles that mediate extracellular communication via transport of various biomolecules such as coding and non-coding RNAs (e.g., miRNA). The exosome communication process impacts several physiological and pathological pathways by modulating protein expression. Recently, investigation into the applicability of exosomal miRNA as diagnostic biomarkers has grown significantly. Here, we evaluated exosomal smRNA, specifically targeting miRNA, and piRNA from the stool samples of IBD model mice as potential markers of IBD.

In the present study, we observed an IBD-like enterocolitis in the IL-10 KO mice (Figure 2). The presence of disease in the IL-10 KO mice was confirmed on clinical signs and histopathology scoring. Significant differences in histopathology severity scores were identified in the large intestine between wild-type control and the IL-10 KO mice. More extensive large intestine damage than small intestine damage falls in line with previous observations (16). However, with a larger sample size, we would have likely also noted a difference in the small intestinal and cecum sections as the data was trending towards showing an increase in scores for the IL-10 KO mice.

When analyzing the scoring data, most mice developed clinical signs of disease and were euthanized by 30 weeks old. In almost all cases, this was due to the rapid weight loss beyond 25% which is considered as a humane endpoint (Figure 1), shortly after developing diarrhea and perianal mucous. However, no mice developed a score of 4, as none of them developed rectal prolapse within the given parameters. It is possible that the IL-10 KO mice do not develop rectal prolapse as part of their disease progression. However, the weight loss may have occurred too rapidly for the disease to progress to prolapse. Of note, all 7 males eventually developed disease, while only 3/7 females developed detectable disease. The differences in disease development rate may be due to a sex bias, but a clear link has yet to be recorded in human cases (26, 27).

The novel fecal smRNA extraction method that we developed produced a robust number of exosomes to use for downstream analysis as shown in both the Nanosight and TapeStation data (Figure 3).

Through analysis of the smRNA data, we identified 504 smRNA that may be implicated in IBD. The time-course analysis using start-, mid-(6 weeks before end-point) and end-point datasets revealed 4 significant smRNA clusters classified based on the trend of expression changes over the course of disease progression (Figure 6). Of note, two clusters, named purple and pink clusters, show sharp expression reduction at the end-point when animals were clinically symptomatic. The pink cluster showed an interesting upward trend in the mid-point, when animals were still clinically asymptomatic. Target gene pathway analysis associated with the pink cluster indicated pro-inflammatory genes are downregulated at the mid-point, when an immediate intervention should be considered although the animals are still asymptomatic. A possible treatment would be administering smRNAs defined in the pink cluster. The expression of the smRNA in the pink cluster is increased at 6 weeks before the end-point (mid-point), however, those expressions then become significantly lower than the start-point. Therefore, continuous administration of those smRNA may prevent the progression of IBD.

As biomarkers of IBD at an earlier stage of pathogenesis, the smRNAs in the pink cluster and the cluster 13 may be combined to determine the disease progression stage.

In summary, our study proposes that fecal exosomal smRNA profiling offers a new groundbreaking opportunity to monitor the inflammatory status of the gut with a capability of detecting its pro-inflammatory (asymptomatic) status. IBD is considered to progress slowly with spatiotemporally dynamic interplay of multiple subsets of immune cells (28). Our next step is to understand from which cells these exosomes are secreted and their physiological impact *in vivo*. In addition, exosomes are also known to contain other biomolecules, such as mRNA, DNA, protein, and lipids. Multi-omic analysis of the fecal exosome may be required as well to understand the comprehensive makeup of the fecal exosome which can then be applied to a development of reengineered exosomes that can be utilized to treat the inflamed colonic lesion (29, 30).

## AUTHOR CONTRIBUTIONS

(I) Conception and design: SM, YIK, (II) Administrative support: NA, (III) Provision of study materials or patients: SM, YIK, (IV) Collection and assembly of data: SM, HMA, YIK, (V) Data analysis and interpretation: SM, ST, HMA, YIK, (VI) Manuscript writing: All authors, (VII) Final approval of manuscript: All authors

## ACKNOWLEDGEMENTS

We thank Georgina Bixler, MS, Robert Brucklacher, MS, Emily Demchak, BS, and Teodora Orendovic, PhD of Penn State College of Medicine Genome Sciences Facility for small RNA-sequencing services. We also thank Han Chen, MD, PhD of Penn State College of Medicine Transmission Electron Microscopy Facility for assistance with transmission electron microscopy images, and Jeffrey M Sundstrom, MD, PhD and Yuanjun Zhao, PhD of Penn State College of Medicine Department of Ophthalmology for assistance with NanoSight data. This work was supported by the Laboratory Animal Medicine MS program at Penn State College of Medicine Department of Comparative Medicine.

## FOOTNOTE

### Reporting Checklist

The authors have completed the ARRIVE reporting checklist.

### Data Sharing Statement

All raw data and processed data of RNA sequencing are available (GSE167239). The datasets generated during the study are available from the corresponding author on reasonable request.

## Ethical Statement

The authors are accountable for all aspects of the work in ensuring that questions related to the accuracy or integrity of any part of the work are appropriately investigated and resolved. All experiments were conducted in accordance with institutional guidelines, the Guide for the Care and Use of Laboratory Animals (Institute for Laboratory Animal Research, 2011) and approved by the Penn State College of Medicine Institutional Animal Care and Uses Committee.

