## Supplementary material for "Dynamic expression of smRNA from fecal exosome in disease progression of an inflammatory bowel disorder mouse model": Supplementary Information Manning et al biorxiv.docx

Yuka Imamura Kawasawa Ph.D.

Penn State University College of Medicine

500 University Dr. Mail Code H093

Hershey, PA 17033

**Supplementary Fig. S1**
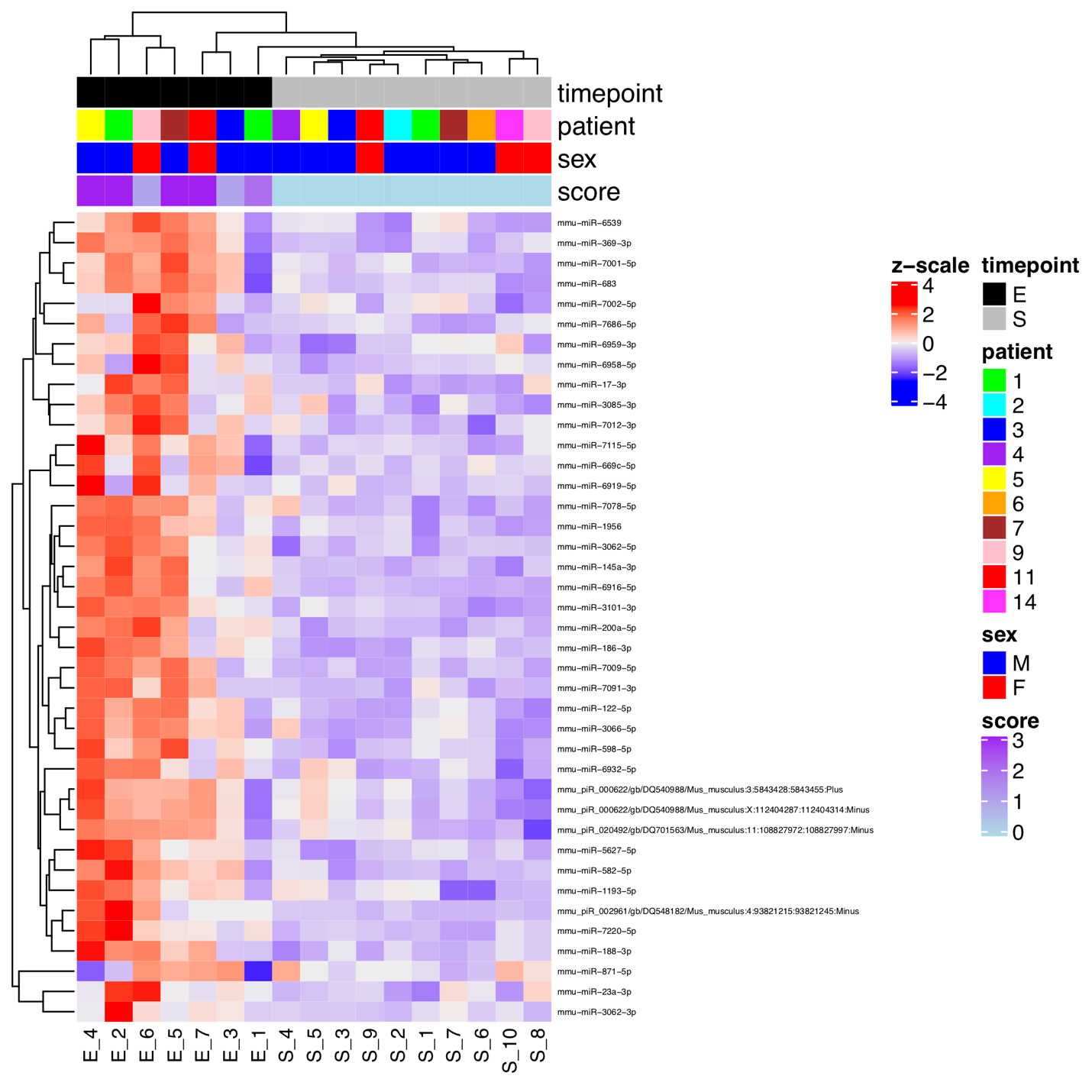

**Supplementary Fig.S1** **Differentially expressed genes between before symptom onset and terminal sedation.** A clustered heatmap of the 40 differentially expressed smRNAs that are consistently called significantly different among 4 algorithms, edgeR, DESeq2, NOI-seq, and limma-voom. The annotation bars depict sample information for its timepoint, mouse identity (patient), sex, and disease scores when feces were sampled, respectively.

**Supplementary Table.S1** **Distinct clusters off differentially expressed smRNAs that manifested time-course dependent changes in IL-10 KO mice.**

| name | logFC by edgeR | adjusted p by edgeR | cluster by STEM |
| --- | --- | --- | --- |
| mmu-miR-138-5p | -4.35 | 2.05E-07 | 0 |
| mmu-miR-448-3p | -2.89 | 3.50E-06 | 0 |
| mmu-miR-543-5p | -3.25 | 1.63E-04 | 0 |
| mmu-miR-7067-3p | -2.77 | 3.50E-04 | 0 |
| mmu-miR-6945-3p | -2.75 | 3.69E-04 | 0 |
| mmu-miR-6899-5p | -3.53 | 4.64E-04 | 0 |
| mmu-miR-1912-3p | -2.13 | 5.47E-04 | 0 |
| mmu-miR-6903-5p | -2.30 | 5.75E-04 | 0 |
| mmu-miR-7011-3p | -3.60 | 7.59E-04 | 0 |
| mmu-miR-192-5p | -2.73 | 1.01E-03 | 0 |
| mmu-miR-185-3p | -2.89 | 1.13E-03 | 0 |
| mmu-miR-770-5p | -1.92 | 1.94E-03 | 0 |
| mmu-miR-7056-3p | -2.92 | 2.14E-03 | 0 |
| mmu-miR-363-3p | -2.13 | 2.39E-03 | 0 |
| mmu-miR-6955-3p | -1.95 | 3.36E-03 | 0 |
| mmu-miR-7689-3p | -3.28 | 5.64E-03 | 0 |
| mmu-miR-3072-5p | -2.36 | 1.11E-02 | 0 |
| mmu-miR-1247-5p | -1.41 | 3.68E-02 | 0 |
| mmu-miR-140-5p | -2.11 | 2.79E-03 | 1 |
| mmu-miR-193b-3p | -1.72 | 9.93E-03 | 1 |
| mmu-miR-130c | -1.64 | 2.00E-02 | 1 |
| mmu-miR-3090-3p | -1.82 | 2.14E-02 | 1 |
| mmu-miR-7689-5p | -1.82 | 2.22E-02 | 1 |
| mmu-miR-20b-5p | -1.40 | 2.29E-02 | 1 |
| mmu-miR-6925-3p | -1.42 | 3.43E-02 | 1 |
| mmu-miR-133a-3p | -1.96 | 3.98E-02 | 1 |
| mmu-miR-3106-3p | -2.53 | 4.00E-02 | 1 |
| mmu-miR-871-5p | 2.89 | 6.94E-05 | blue |
| mmu-miR-6370 | -1.28 | 1.45E-02 | blue |
| mmu-miR-3101-5p | 2.40 | 1.46E-02 | blue |
| mmu-miR-363-5p | -1.22 | 1.49E-02 | blue |
| mmu-miR-547-5p | 2.03 | 2.39E-02 | blue |
| mmu-miR-7220-3p | -1.17 | 2.55E-02 | blue |
| mmu-miR-211-5p | 1.35 | 3.28E-02 | blue |
| mmu-miR-431-3p | 1.25 | 3.73E-02 | blue |
| mmu-miR-741-3p | -1.49 | 3.76E-02 | blue |
| mmu-miR-497a-5p | 1.53 | 4.23E-02 | blue |
| mmu-miR-5128 | -1.30 | 4.43E-02 | blue |
| mmu-miR-423-3p | -2.56 | 4.50E-02 | blue |
| mmu-miR-1193-5p | 4.26 | 3.35E-09 | 11 |
| mmu-miR-582-5p | 4.80 | 2.64E-07 | 11 |
| mmu-miR-6958-5p | 3.66 | 8.21E-07 | 11 |
| mmu-miR-6919-5p | 3.50 | 2.09E-06 | 11 |
| mmu-miR-294-3p | 3.34 | 2.54E-06 | 11 |
| mmu-miR-7000-5p | 3.20 | 2.54E-06 | 11 |
| mmu-miR-7001-5p | 3.29 | 7.47E-06 | 11 |
| mmu-miR-1955-3p | 2.95 | 1.45E-05 | 11 |
| mmu-miR-323-3p | 4.03 | 1.76E-05 | 11 |
| mmu-miR-3066-5p | 4.01 | 3.08E-05 | 11 |
| mmu-miR-7671-5p | 3.00 | 4.62E-05 | 11 |
| mmu-miR-151-3p | 2.82 | 6.54E-05 | 11 |
| mmu-miR-6959-3p | 2.80 | 6.96E-05 | 11 |
| mmu-miR-429-5p | 3.01 | 9.64E-05 | 11 |
| mmu-miR-27a-5p | 4.41 | 1.16E-04 | 11 |
| mmu-miR-6998-3p | 3.96 | 1.49E-04 | 11 |
| mmu-miR-3079-3p | 2.05 | 2.20E-04 | 11 |
| mmu-miR-3964 | 2.35 | 3.51E-04 | 11 |
| mmu-miR-290a-3p | 2.36 | 6.45E-04 | 11 |
| mmu-miR-6919-3p | 3.45 | 6.80E-04 | 11 |
| mmu-miR-3070-3p | 2.64 | 7.86E-04 | 11 |
| mmu-miR-3473c | 1.58 | 9.12E-04 | 11 |
| mmu-miR-1981-3p | 2.27 | 1.67E-03 | 11 |
| mmu-miR-1932 | 2.18 | 1.87E-03 | 11 |
| mmu-miR-181c-3p | 1.86 | 2.01E-03 | 11 |
| mmu-miR-1963 | 3.11 | 2.11E-03 | 11 |
| mmu-miR-374c-3p | 3.08 | 2.20E-03 | 11 |
| mmu-miR-6941-3p | 2.00 | 3.84E-03 | 11 |
| mmu-miR-184-5p | 2.49 | 4.49E-03 | 11 |
| mmu-miR-292a-3p | 1.77 | 5.69E-03 | 11 |
| mmu-miR-3110-3p | 1.86 | 6.63E-03 | 11 |
| mmu-miR-872-3p | 1.79 | 8.04E-03 | 11 |
| mmu-miR-466d-5p | 2.21 | 8.54E-03 | 11 |
| mmu-miR-7057-5p | 2.03 | 8.86E-03 | 11 |
| mmu-miR-1188-5p | 1.43 | 1.28E-02 | 11 |
| mmu-miR-6418-5p | 1.57 | 2.17E-02 | 11 |
| mmu-miR-6926-3p | 3.04 | 2.17E-02 | 11 |
| mmu-miR-101b-5p | 1.61 | 2.92E-02 | 11 |
| mmu-miR-130b-3p | 1.36 | 3.20E-02 | 11 |
| mmu-miR-8096 | 1.58 | 3.95E-02 | 11 |
| mmu-miR-7661-3p | 1.15 | 4.19E-02 | 11 |
| mmu-miR-6926-5p | 1.30 | 4.59E-02 | 11 |
| mmu-miR-6916-5p | 8.95 | 1.16E-16 | 12 |
| mmu-miR-122-5p | 5.37 | 9.13E-08 | 12 |
| mmu-miR-3062-3p | 6.92 | 2.05E-07 | 12 |
| mmu_piR_020492/gb/DQ701563/Mus_musculus:11:108827972:108827997:Minus | 3.72 | 5.90E-06 | 12 |
| mmu-miR-7079-5p | 3.70 | 9.29E-06 | 12 |
| mmu-miR-6378 | 2.63 | 1.16E-05 | 12 |
| mmu-miR-7677-3p | 4.95 | 1.16E-05 | 12 |
| mmu-miR-8118 | 5.33 | 1.62E-05 | 12 |
| mmu-miR-24-1-5p | 4.57 | 3.04E-05 | 12 |
| mmu-miR-758-3p | 2.82 | 3.37E-05 | 12 |
| mmu-miR-32-5p | 5.57 | 5.58E-05 | 12 |
| mmu-miR-6383 | 2.24 | 2.67E-04 | 12 |
| mmu-miR-143-3p | 2.72 | 3.92E-04 | 12 |
| mmu-miR-141-3p | 2.96 | 4.26E-04 | 12 |
| mmu-miR-7074-3p | 2.32 | 1.02E-03 | 12 |
| mmu-miR-6982-5p | 2.09 | 1.53E-03 | 12 |
| mmu-miR-3059-3p | 2.39 | 1.95E-03 | 12 |
| mmu-miR-7219-3p | 2.45 | 2.10E-03 | 12 |
| mmu-miR-215-5p | 2.70 | 2.39E-03 | 12 |
| mmu-miR-6922-3p | 2.68 | 2.64E-03 | 12 |
| mmu-miR-7090-5p | 1.67 | 3.76E-02 | 12 |
| mmu-miR-145a-3p | 6.11 | 7.28E-13 | 13 |
| mmu-miR-7686-5p | 7.39 | 2.20E-09 | 13 |
| mmu-miR-7220-5p | 8.02 | 2.40E-09 | 13 |
| mmu-miR-1193-3p | 6.68 | 2.11E-08 | 13 |
| mmu_piR_002961/gb/DQ548182/Mus_musculus:4:93821215:93821245:Minus | 11.65 | 3.63E-08 | 13 |
| mmu-miR-186-3p | 5.05 | 2.91E-07 | 13 |
| mmu-miR-17-3p | 6.14 | 2.44E-06 | 13 |
| mmu-miR-3062-5p | 3.82 | 2.54E-06 | 13 |
| mmu-miR-8115 | 2.83 | 5.09E-06 | 13 |
| mmu-miR-5123 | 3.08 | 8.62E-06 | 13 |
| mmu-miR-3085-3p | 4.28 | 1.03E-05 | 13 |
| mmu-miR-152-5p | 2.36 | 1.29E-05 | 13 |
| mmu-miR-122-3p | 4.44 | 1.77E-05 | 13 |
| mmu_piR_000622/gb/DQ540988/Mus_musculus:X:112404287:112404314:Minus | 3.54 | 4.38E-05 | 13 |
| mmu-miR-7094-3p | 5.70 | 2.27E-04 | 13 |
| mmu-miR-874-5p | 2.62 | 7.72E-04 | 13 |
| mmu-miR-666-3p | 2.17 | 8.90E-04 | 13 |
| mmu-miR-6914-5p | 3.10 | 1.01E-03 | 13 |
| mmu-miR-7671-3p | 2.22 | 2.44E-03 | 13 |
| mmu-miR-3068-5p | 1.90 | 2.64E-03 | 13 |
| mmu-miR-378a-5p | 3.13 | 2.71E-03 | 13 |
| mmu-miR-6963-5p | 1.93 | 1.88E-02 | 13 |
| mmu-miR-1298-5p | 1.66 | 2.99E-02 | 13 |
| mmu-miR-701-3p | 1.83 | 4.29E-02 | 13 |
| mmu-miR-126b-5p | 1.44 | 4.51E-02 | 13 |
| mmu-miR-465d-5p | 5.85 | 4.61E-06 | green |
| mmu-miR-18b-3p | 3.32 | 6.79E-06 | green |
| mmu-miR-669h-5p | 4.18 | 4.89E-04 | green |
| mmu-miR-497b | 3.51 | 4.99E-04 | green |
| mmu-miR-1898 | 2.82 | 5.65E-04 | green |
| mmu-miR-7219-5p | 2.78 | 5.65E-04 | green |
| mmu-miR-873b | 2.85 | 7.60E-04 | green |
| mmu-miR-708-5p | 2.88 | 8.14E-04 | green |
| mmu-miR-875-3p | 2.03 | 1.07E-03 | green |
| mmu-miR-28c | 3.37 | 1.07E-03 | green |
| mmu-miR-199a-5p | 2.31 | 1.52E-03 | green |
| mmu-miR-30e-3p | 3.22 | 1.75E-03 | green |
| mmu-miR-98-3p | 1.83 | 1.95E-03 | green |
| mmu-miR-291a-3p | 2.54 | 2.68E-03 | green |
| mmu-miR-344-5p | 3.57 | 4.06E-03 | green |
| mmu-miR-1231-5p | 2.25 | 4.35E-03 | green |
| mmu-miR-540-3p | 1.72 | 6.65E-03 | green |
| mmu-miR-7646-5p | 1.48 | 7.01E-03 | green |
| mmu-miR-196b-3p | 2.16 | 7.43E-03 | green |
| mmu-miR-409-3p | 1.76 | 7.78E-03 | green |
| mmu-miR-467b-5p | 2.33 | 7.99E-03 | green |
| mmu-miR-7212-5p | 1.91 | 8.79E-03 | green |
| mmu-miR-546 | 1.86 | 1.29E-02 | green |
| mmu-miR-3102-3p | 1.54 | 1.42E-02 | green |
| mmu-miR-1941-3p | 1.30 | 1.91E-02 | green |
| mmu-miR-714 | 1.61 | 2.07E-02 | green |
| mmu-miR-185-5p | 1.98 | 2.09E-02 | green |
| mmu-miR-6970-3p | 1.46 | 2.09E-02 | green |
| mmu-miR-6412 | 1.52 | 2.15E-02 | green |
| mmu-miR-382-5p | 1.45 | 2.22E-02 | green |
| mmu-miR-6901-3p | 1.57 | 2.35E-02 | green |
| mmu-miR-7006-3p | 2.69 | 2.61E-02 | green |
| mmu-miR-6908-5p | 1.50 | 3.15E-02 | green |
| mmu-miR-7679-3p | 1.79 | 3.54E-02 | green |
| mmu-miR-490-5p | 1.50 | 4.12E-02 | green |
| mmu-miR-3569-5p | 1.08 | 4.23E-02 | green |
| mmu-miR-6948-5p | 1.22 | 4.23E-02 | green |
| mmu-miR-433-3p | 1.26 | 4.50E-02 | green |
| mmu-miR-6952-5p | 1.18 | 4.53E-02 | green |
| mmu-miR-879-5p | 1.38 | 4.53E-02 | green |
| mmu-miR-181b-5p | 1.35 | 4.82E-02 | green |
| mmu-miR-7012-3p | 4.13 | 7.47E-10 | 15 |
| mmu-miR-3101-3p | 4.93 | 1.33E-09 | 15 |
| mmu-miR-669c-5p | 4.42 | 3.50E-09 | 15 |
| mmu-miR-598-5p | 3.76 | 7.37E-09 | 15 |
| mmu-miR-683 | 3.80 | 4.44E-07 | 15 |
| mmu-miR-7091-3p | 5.38 | 4.80E-07 | 15 |
| mmu-miR-501-5p | 3.53 | 4.62E-06 | 15 |
| mmu-miR-7020-3p | 2.55 | 1.76E-05 | 15 |
| mmu-miR-23a-3p | 3.25 | 1.16E-04 | 15 |
| mmu_piR_000691/gb/DQ541218/Mus_musculus:8:126494409:126494434:Minus | 2.98 | 2.13E-04 | 15 |
| mmu-miR-6898-5p | 3.80 | 2.41E-04 | 15 |
| mmu-miR-223-3p | 2.49 | 3.61E-04 | 15 |
| mmu-miR-6932-5p | 2.68 | 6.63E-04 | 15 |
| mmu-miR-5129-3p | 3.69 | 1.48E-03 | 15 |
| mmu-miR-1188-3p | 1.98 | 1.61E-03 | 15 |
| mmu-miR-125b-2-3p | 2.29 | 2.05E-03 | 15 |
| mmu-miR-7094b-2-5p | 2.21 | 1.70E-02 | 15 |
| mmu-miR-7215-3p | 1.46 | 1.87E-02 | 15 |
| mmu-miR-452-3p | 1.78 | 2.40E-02 | 15 |
| mmu-miR-3057-5p | 1.34 | 3.13E-02 | 15 |
| mmu-miR-125a-3p | 1.56 | 3.76E-02 | 15 |
| mmu-miR-742-3p | 1.41 | 4.23E-02 | 15 |
| mmu-miR-7066-3p | -4.25 | 5.54E-08 | 2 |
| mmu-miR-6943-3p | -3.74 | 2.41E-05 | 2 |
| mmu-miR-6897-5p | -3.01 | 1.62E-03 | 2 |
| mmu-miR-6419 | -2.83 | 2.25E-03 | 2 |
| mmu-miR-3074-2-3p | -2.88 | 2.28E-03 | 2 |
| mmu-miR-3091-3p | -2.51 | 3.53E-03 | 2 |
| mmu-miR-6392-3p | -1.99 | 3.76E-03 | 2 |
| mmu-miR-322-3p | -2.30 | 4.49E-03 | 2 |
| mmu-miR-3092-3p | -1.67 | 4.62E-03 | 2 |
| mmu-miR-19b-1-5p | -1.70 | 1.13E-02 | 2 |
| mmu-miR-5119 | 2.24 | 1.31E-02 | 2 |
| mmu-miR-7031-5p | -1.87 | 2.07E-02 | 2 |
| mmu-miR-1249-3p | -1.28 | 3.10E-02 | 2 |
| mmu-miR-7058-5p | -2.09 | 4.50E-02 | 2 |
| mmu-miR-7042-3p | -3.46 | 6.67E-07 | 3 |
| mmu-miR-3544-5p | -5.56 | 1.18E-04 | 3 |
| mmu-miR-653-3p | -3.30 | 1.34E-04 | 3 |
| mmu-miR-7024-3p | -2.90 | 3.02E-04 | 3 |
| mmu_piR_002738/gb/DQ547526/Mus_musculus:1:57297924:57297953:Minus | -4.01 | 3.59E-04 | 3 |
| mmu-miR-3073a-3p | -3.92 | 5.53E-04 | 3 |
| mmu-miR-3070-5p | -2.70 | 6.63E-04 | 3 |
| mmu-miR-7028-5p | -2.82 | 7.55E-04 | 3 |
| mmu-miR-335-3p | -4.16 | 9.03E-04 | 3 |
| mmu_piR_002962/gb/DQ548183/Mus_musculus:5:146565277:146565304:Plus | -4.54 | 1.95E-03 | 3 |
| mmu-miR-7010-5p | -1.71 | 4.06E-03 | 3 |
| mmu-miR-218-2-3p | -1.59 | 4.57E-03 | 3 |
| mmu-miR-7018-5p | -2.47 | 4.57E-03 | 3 |
| mmu-miR-7240-5p | -2.51 | 8.61E-03 | 3 |
| mmu-miR-2861 | -1.83 | 9.27E-03 | 3 |
| mmu-miR-7077-3p | -3.22 | 1.54E-06 | 4 |
| mmu-miR-7680-3p | -3.63 | 5.90E-06 | 4 |
| mmu-miR-744-5p | -2.79 | 1.03E-05 | 4 |
| mmu-miR-7665-5p | -2.73 | 3.56E-05 | 4 |
| mmu-miR-6997-5p | -2.82 | 1.16E-04 | 4 |
| mmu-miR-1948-5p | -2.92 | 2.22E-04 | 4 |
| mmu-miR-204-3p | -1.94 | 1.42E-03 | 4 |
| mmu-miR-1934-5p | -2.43 | 1.91E-03 | 4 |
| mmu-miR-208a-5p | -3.02 | 2.71E-03 | 4 |
| mmu-miR-6394 | -1.99 | 5.78E-03 | 4 |
| mmu-miR-6979-3p | -2.73 | 8.84E-03 | 4 |
| mmu-miR-5107-3p | -2.10 | 1.02E-02 | 4 |
| mmu-miR-673-3p | -2.20 | 1.30E-02 | 4 |
| mmu-miR-375-3p | -2.20 | 2.39E-02 | 4 |
| mmu-miR-7075-5p | -1.46 | 2.73E-02 | 4 |
| mmu-miR-7018-3p | -1.36 | 4.25E-02 | 4 |
| mmu-miR-93-3p | 2.90 | 8.15E-05 | 5 |
| mmu-miR-1195 | 1.61 | 5.09E-03 | 5 |
| mmu-miR-466q | 1.99 | 2.05E-02 | 5 |
| mmu-miR-200a-5p | 3.61 | 1.94E-10 | 6 |
| mmu-miR-7115-5p | 4.89 | 3.76E-10 | 6 |
| mmu-miR-6539 | 3.56 | 2.54E-06 | 6 |
| mmu-miR-7677-5p | 9.58 | 3.40E-06 | 6 |
| mmu-miR-7071-3p | 3.35 | 2.94E-05 | 6 |
| mmu-miR-383-5p | 1.94 | 6.31E-05 | 6 |
| mmu-miR-5627-3p | 2.25 | 6.10E-04 | 6 |
| mmu-miR-674-5p | 2.87 | 7.55E-04 | 6 |
| mmu-miR-191-5p | 1.93 | 7.56E-04 | 6 |
| mmu-miR-7082-5p | 2.13 | 1.72E-03 | 6 |
| mmu-miR-7002-5p | 2.01 | 2.02E-03 | 6 |
| mmu-miR-6993-5p | 2.94 | 4.75E-03 | 6 |
| mmu-miR-8101 | 1.78 | 8.98E-03 | 6 |
| mmu-miR-8110 | 2.10 | 1.13E-02 | 6 |
| mmu-miR-1966-3p | 2.08 | 1.21E-02 | 6 |
| mmu-miR-344g-3p | 1.58 | 1.26E-02 | 6 |
| mmu-miR-7011-5p | 1.45 | 3.07E-02 | 6 |
| mmu-miR-682 | -4.07 | 2.05E-07 | pink |
| mmu-miR-672-3p | -2.98 | 2.31E-06 | pink |
| mmu-miR-7084-3p | -3.90 | 2.37E-06 | pink |
| mmu-miR-6415 | -3.89 | 2.54E-06 | pink |
| mmu-miR-6960-3p | -3.22 | 3.56E-05 | pink |
| mmu-miR-677-5p | -3.03 | 1.11E-04 | pink |
| mmu-miR-7118-3p | -2.67 | 4.10E-04 | pink |
| mmu-miR-367-5p | -2.85 | 5.45E-04 | pink |
| mmu-miR-7007-5p | -1.96 | 7.26E-04 | pink |
| mmu-miR-7238-3p | -2.90 | 1.07E-03 | pink |
| mmu-miR-5618-5p | -2.49 | 1.10E-03 | pink |
| mmu-miR-6952-3p | -2.89 | 1.13E-03 | pink |
| mmu-miR-1964-5p | -3.17 | 1.25E-03 | pink |
| mmu-miR-1843b-3p | -1.87 | 1.88E-03 | pink |
| mmu-miR-687 | -1.80 | 2.66E-03 | pink |
| mmu-miR-431-5p | -3.38 | 2.86E-03 | pink |
| mmu-miR-509-3p | -1.62 | 3.41E-03 | pink |
| mmu-miR-675-5p | -1.67 | 3.60E-03 | pink |
| mmu-miR-6983-5p | -1.82 | 3.61E-03 | pink |
| mmu-miR-6977-3p | -1.85 | 4.06E-03 | pink |
| mmu-miR-7217-5p | 3.24 | 4.11E-03 | pink |
| mmu-miR-6377 | -2.25 | 4.27E-03 | pink |
| mmu-miR-1927 | -1.73 | 5.52E-03 | pink |
| mmu-miR-6994-5p | -2.58 | 5.52E-03 | pink |
| mmu-miR-7664-5p | -2.51 | 5.73E-03 | pink |
| mmu-miR-99b-3p | -1.77 | 6.28E-03 | pink |
| mmu-miR-6998-5p | -1.68 | 6.29E-03 | pink |
| mmu-miR-1948-3p | -2.11 | 6.63E-03 | pink |
| mmu-miR-6346 | -1.77 | 7.43E-03 | pink |
| mmu-miR-7049-3p | -1.71 | 7.48E-03 | pink |
| mmu-miR-6386 | -1.94 | 8.92E-03 | pink |
| mmu-let-7c-1-3p | -2.78 | 9.25E-03 | pink |
| mmu-miR-367-3p | -1.58 | 9.80E-03 | pink |
| mmu-miR-6238 | -1.80 | 1.23E-02 | pink |
| mmu-miR-5617-5p | -1.48 | 1.36E-02 | pink |
| mmu-miR-6961-5p | -1.74 | 2.14E-02 | pink |
| mmu-miR-532-3p | -2.09 | 2.32E-02 | pink |
| mmu-miR-7004-5p | -1.48 | 3.07E-02 | pink |
| mmu-miR-3061-5p | -2.23 | 3.20E-02 | pink |
| mmu-miR-5133 | -1.85 | 3.37E-02 | pink |
| mmu-miR-106b-3p | -1.19 | 4.19E-02 | pink |
| mmu-miR-342-3p | -1.30 | 4.59E-02 | pink |
| mmu-miR-1956 | 6.66 | 1.37E-11 | 8 |
| mmu-miR-7009-5p | 4.52 | 4.33E-11 | 8 |
| mmu-miR-188-3p | 4.48 | 5.79E-10 | 8 |
| mmu-miR-369-3p | 2.96 | 1.52E-08 | 8 |
| mmu-miR-5627-5p | 3.73 | 8.64E-08 | 8 |
| mmu-miR-7078-5p | 5.92 | 4.41E-07 | 8 |
| mmu-miR-6912-5p | 3.15 | 4.80E-07 | 8 |
| mmu-miR-6953-3p | 4.95 | 4.95E-06 | 8 |
| mmu-miR-5114 | 5.11 | 5.09E-06 | 8 |
| mmu-miR-3077-3p | 3.25 | 1.12E-05 | 8 |
| mmu-miR-411-3p | 3.98 | 3.56E-05 | 8 |
| mmu_piR_000622/gb/DQ540988/Mus_musculus:3:5843428:5843455:Plus | 3.59 | 9.98E-05 | 8 |
| mmu-miR-130a-3p | 3.63 | 1.16E-04 | 8 |
| mmu-miR-7038-5p | 3.89 | 1.32E-04 | 8 |
| mmu-miR-6356 | 3.15 | 1.83E-04 | 8 |
| mmu-miR-6915-3p | 2.76 | 2.67E-04 | 8 |
| mmu-miR-679-3p | 3.39 | 2.91E-04 | 8 |
| mmu-miR-7017-5p | 2.40 | 3.01E-04 | 8 |
| mmu-miR-101b-3p | 5.22 | 4.40E-04 | 8 |
| mmu-miR-7063-3p | 4.85 | 4.64E-04 | 8 |
| mmu-miR-3065-5p | 2.51 | 4.91E-04 | 8 |
| mmu-miR-3082-5p | 2.36 | 4.91E-04 | 8 |
| mmu-miR-505-5p | 2.95 | 6.27E-04 | 8 |
| mmu-miR-1899 | 3.90 | 1.48E-03 | 8 |
| mmu-miR-344d-2-5p | 2.72 | 2.01E-03 | 8 |
| mmu-miR-1900 | 1.91 | 3.14E-03 | 8 |
| mmu-miR-9769-5p | 1.91 | 4.03E-03 | 8 |
| mmu-miR-7657-5p | 2.41 | 5.32E-03 | 8 |
| mmu-miR-7211-3p | 1.64 | 5.75E-03 | 8 |
| mmu-miR-344c-3p | 1.43 | 6.23E-03 | 8 |
| mmu-miR-200b-3p | 3.46 | 8.07E-03 | 8 |
| mmu-miR-7213-3p | 2.05 | 1.03E-02 | 8 |
| mmu_piR_000622/gb/DQ540988/Mus_musculus:2:5296560:5296587:Minus | 2.05 | 1.08E-02 | 8 |
| mmu-miR-697 | 1.35 | 1.31E-02 | 8 |
| mmu-miR-3474 | 1.96 | 1.34E-02 | 8 |
| mmu-miR-6368 | 1.76 | 1.51E-02 | 8 |
| mmu-miR-3095-3p | 1.60 | 1.67E-02 | 8 |
| mmu-miR-137-3p | 1.50 | 1.78E-02 | 8 |
| mmu-miR-483-3p | 1.41 | 2.05E-02 | 8 |
| mmu-miR-331-3p | 1.49 | 2.39E-02 | 8 |
| mmu-miR-6358 | 1.59 | 3.24E-02 | 8 |
| mmu-miR-1951 | 1.66 | 3.28E-02 | 8 |
| mmu-miR-1897-3p | 1.60 | 3.87E-02 | 8 |
| mmu-miR-6416-3p | 1.37 | 4.59E-02 | 8 |
| mmu-miR-708-3p | -2.30 | 1.16E-04 | purple |
| mmu-miR-370-5p | -2.94 | 1.43E-04 | purple |
| mmu-miR-7030-5p | -2.34 | 6.54E-04 | purple |
| mmu-miR-3552 | -2.95 | 7.59E-04 | purple |
| mmu-miR-7074-5p | -2.41 | 1.48E-03 | purple |
| mmu-miR-1969 | -3.27 | 1.72E-03 | purple |
| mmu-miR-344i | -2.10 | 1.75E-03 | purple |
| mmu-miR-362-3p | -2.02 | 1.75E-03 | purple |
| mmu-miR-208a-3p | -1.67 | 1.81E-03 | purple |
| mmu-miR-1962 | -1.67 | 2.05E-03 | purple |
| mmu-miR-7052-5p | -1.62 | 2.05E-03 | purple |
| mmu-miR-499-5p | -2.73 | 2.10E-03 | purple |
| mmu-miR-7682-3p | -2.50 | 2.64E-03 | purple |
| mmu-miR-8095 | -1.83 | 3.31E-03 | purple |
| mmu-miR-22-5p | -1.75 | 3.32E-03 | purple |
| mmu-miR-6897-3p | -1.79 | 3.40E-03 | purple |
| mmu-miR-1957b | -1.78 | 3.58E-03 | purple |
| mmu-miR-30c-2-3p | -2.26 | 3.76E-03 | purple |
| mmu-miR-8117 | -1.84 | 4.57E-03 | purple |
| mmu-miR-3475-3p | -2.91 | 5.18E-03 | purple |
| mmu-miR-7211-5p | -1.61 | 5.73E-03 | purple |
| mmu-miR-7031-3p | -1.80 | 6.08E-03 | purple |
| mmu-miR-3067-3p | -2.18 | 6.23E-03 | purple |
| mmu-miR-7674-3p | -1.93 | 6.40E-03 | purple |
| mmu-miR-1983 | -1.76 | 7.19E-03 | purple |
| mmu-miR-377-5p | -3.70 | 7.51E-03 | purple |
| mmu-miR-8093 | -1.48 | 8.98E-03 | purple |
| mmu-miR-7033-5p | -1.45 | 9.51E-03 | purple |
| mmu-miR-743b-5p | -1.59 | 9.93E-03 | purple |
| mmu-miR-494-3p | -1.57 | 1.02E-02 | purple |
| mmu-miR-6965-5p | -1.56 | 1.02E-02 | purple |
| mmu-miR-7649-3p | -1.55 | 1.14E-02 | purple |
| mmu-miR-7225-5p | -2.73 | 1.18E-02 | purple |
| mmu-miR-7119-5p | -2.28 | 1.21E-02 | purple |
| mmu-miR-3569-3p | -2.76 | 1.27E-02 | purple |
| mmu-miR-1934-3p | -1.62 | 1.30E-02 | purple |
| mmu-miR-6946-5p | -1.63 | 1.30E-02 | purple |
| mmu-miR-3083-3p | -1.68 | 1.31E-02 | purple |
| mmu-miR-31-5p | -1.78 | 1.39E-02 | purple |
| mmu-miR-6936-5p | -1.37 | 1.39E-02 | purple |
| mmu-miR-7024-5p | -2.00 | 1.52E-02 | purple |
| mmu-miR-1946a | -2.55 | 1.53E-02 | purple |
| mmu-miR-187-5p | -2.12 | 1.72E-02 | purple |
| mmu-miR-7040-5p | -1.43 | 1.78E-02 | purple |
| mmu-miR-6921-3p | -1.47 | 1.92E-02 | purple |
| mmu_piR_001538/gb/DQ543617/Mus_musculus:6:87920828:87920858:Minus | -1.78 | 1.95E-02 | purple |
| mmu-miR-194-5p | -2.03 | 2.42E-02 | purple |
| mmu-miR-7681-5p | -1.13 | 2.42E-02 | purple |
| mmu-miR-6959-5p | -1.40 | 2.51E-02 | purple |
| mmu-miR-6351 | -1.23 | 2.52E-02 | purple |
| mmu-miR-466l-3p | -1.70 | 2.61E-02 | purple |
| mmu-miR-3154 | -1.90 | 2.61E-02 | purple |
| mmu-miR-34b-3p | -1.90 | 2.66E-02 | purple |
| mmu-miR-486a-5p | -1.64 | 2.67E-02 | purple |
| mmu-miR-671-5p | -1.06 | 2.72E-02 | purple |
| mmu-miR-19a-3p | -1.31 | 3.11E-02 | purple |
| mmu-miR-678 | -2.21 | 3.11E-02 | purple |
| mmu-miR-6939-5p | -1.31 | 3.16E-02 | purple |
| mmu-miR-3081-5p | -1.27 | 3.24E-02 | purple |
| mmu-miR-3089-3p | -1.47 | 3.41E-02 | purple |
| mmu-miR-3473d | -1.55 | 3.59E-02 | purple |
| mmu-miR-106a-5p | -1.83 | 3.74E-02 | purple |
| mmu-miR-344f-3p | -1.31 | 3.76E-02 | purple |
| mmu-miR-434-3p | -1.20 | 4.13E-02 | purple |
| mmu-miR-7646-3p | -1.41 | 4.19E-02 | purple |
| mmu-miR-1198-5p | -1.14 | 4.23E-02 | purple |
| mmu-miR-421-3p | -1.72 | 4.23E-02 | purple |
| mmu-miR-1191b-5p | -1.32 | 4.50E-02 | purple |
| mmu-miR-1957a | -1.33 | 4.50E-02 | purple |
| mmu-miR-324-3p | -1.85 | 4.50E-02 | purple |
| mmu-miR-6410 | -1.43 | 4.74E-02 | purple |
| mmu-miR-7053-3p | -1.35 | 4.93E-02 | purple |
| mmu-miR-6347 | -1.95 | 4.94E-02 | purple |

A total 414 smRNAs are listed, while a total of 504 smRNAs were applied to the STEM analysis. 90 smRNAs showed inconsistent temporal changes among samples, and thus are filtered out in the early quality control step of the STEM analysis.
